## Supplementary Material for "Dynamic persistence of intracellular bacterial communities of uropathogenic *Escherichia coli* in a human bladder-chip model of urinary tract infections"

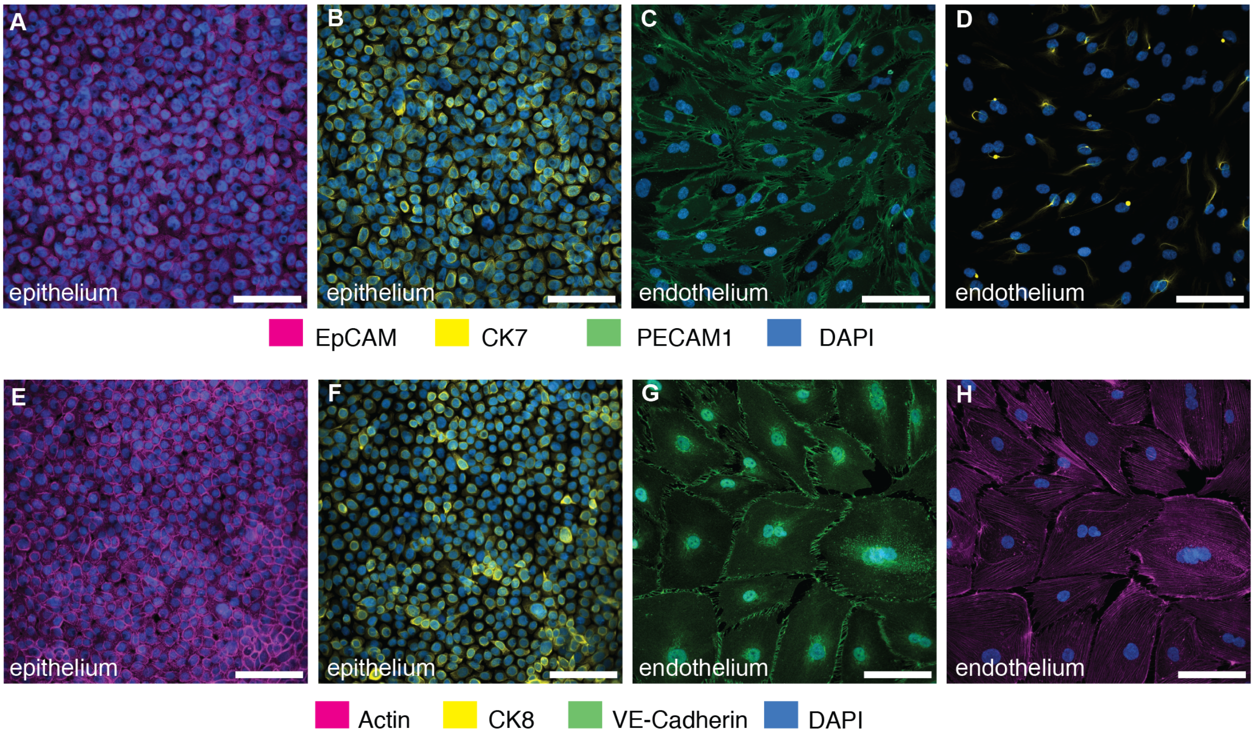


**Figure S1: Characterization of co-cultures of bladder epithelial cells and bladder endothelial cells in bladder-chip.**

Immunofluorescence characterization of HTB9 bladder epithelial cells in bladder-chip for Epithelial Cell Adhesion Molecule (EpCAM) (**A**), cytokeratin 7 (CK7) (**B**), and cytokeratin 8 (CK8), a marker for differentiated urothelial cells (**F**). Immunofluorescence characterization of primary human bladder microvascular endothelial cells in bladder-chip for tight junction markers such as Platelet Endothelial Cell Adhesion Molecule-1 (PECAM-1) (**C**) and vascular endothelial cadherin (VE-cadherin) (**G**). Some endothelial cells also express CK7 (**D**). Filamentous actin staining for epithelial (**E**) and endothelial (**H**) cells. Cell nuclei were labeled with DAPI (azure) in all panels. Scale bars, 50 µm in all panels.

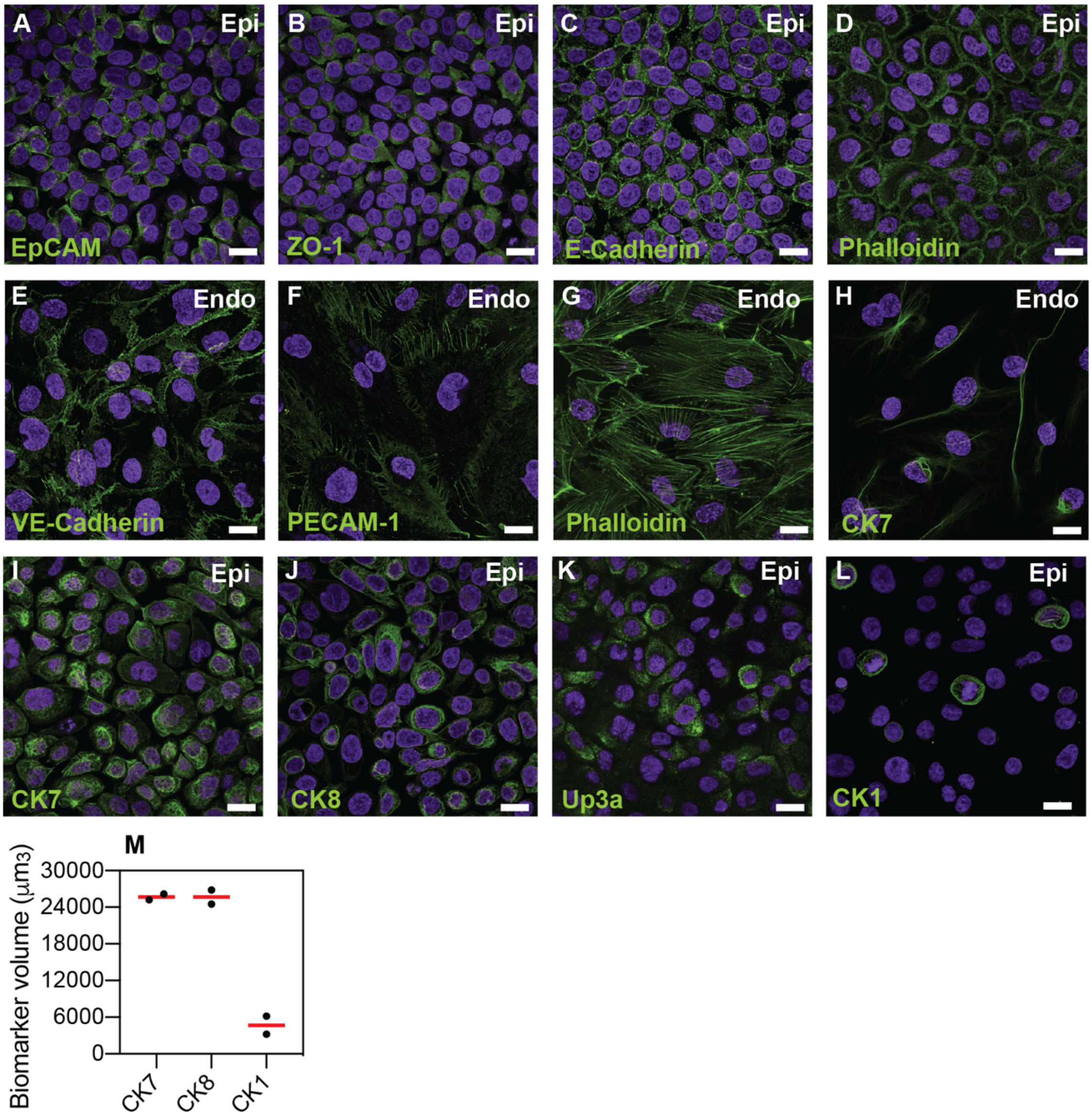


**Figure S2: Characterization of monocultures of HTB9 bladder epithelial cells and HMVEC-Bd bladder microvascular endothelial cells.**
Characterization of the HTB9 bladder epithelial cells for epithelial tight junction markers such as Epithelial Cell Adhesion Molecule (EpCAM) (**A**), epithelial cadherins (E-cadherin) (**B**), Zonula Occludens-1 (ZO-1) (**C**), and filamentous actin (Phalloidin) (**D**). Bladder endothelial cells express tight junction markers such as vascular, Platelet Endothelial Cell Adhesion Molecule-1 (PECAM-1) (**E**), endothelial cadherin (VE-cadherin) (**F**) and filamentous actin (Phalloidin) (**G**). Some endothelial cells also showed staining for CK7 (**H**). Characterization of the HTB9 bladder epithelial cell line for the uroepithelial cell marker cytokeratin 7 (CK7) (**I**), for umbrella cell specific markers cytokeratin 8 (CK8) (**J**) and uroplakin 3a (Up3a) (**K**), and the basal cell marker cytokeratin 1 (CK1) (**L**). CK1 expression was sparse and lower than CK7 and CK8, data obtained from 2 fields of view in an ibidi µ-Slide 8 well (**M**). Red lines represent the median value. Epithelial and endothelial cells were grown to ca. 75-90% confluence in ibidi 8-wells. Cell nuclei were labeled with DAPI (cyan). Scale bars, 10 µm in (**A-L**).

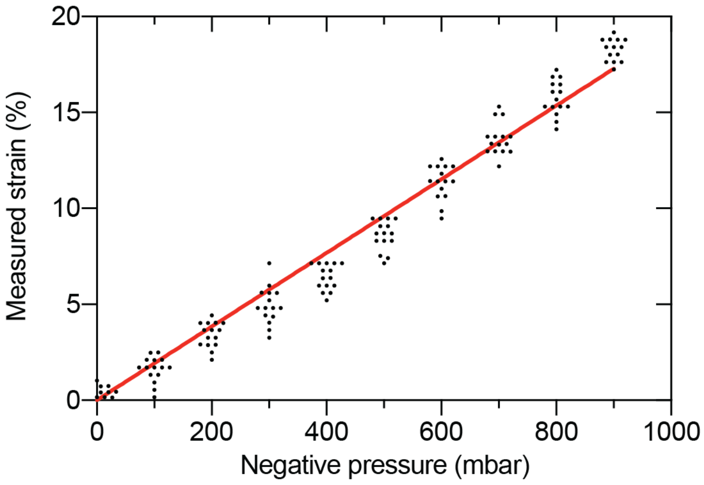


**Figure S3: Quantification of the linear strain in the PDMS membrane as a function of applied negative pressure in the vacuum channels of the bladder-chip.**
Pore-to-pore distance was measured in the PDMS membrane (n=14) on human bladder chip under different values of applied pressure and used to calculate the linear strain ($\Delta l = \frac{l_s - l_r}{l_r}$). $l_s$ and $l_r$ refer to the pore-to-pore distance in the stretched ($l_s$) and relaxed state ($l_r$).

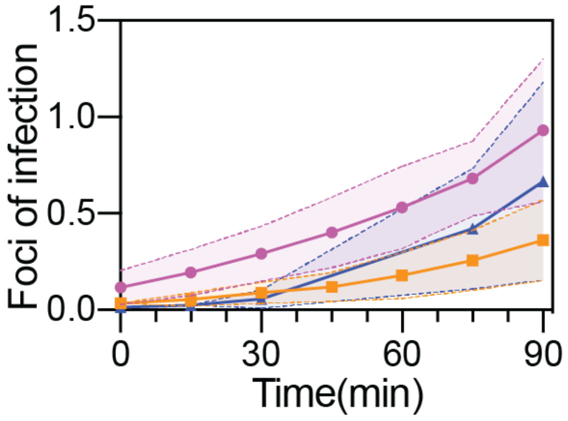


**Figure S4: Quantification of UPEC attachment to bladder epithelial cells on-chip under flow.**

Ratio of number of attached UPEC to the average number of epithelial cells in n=25 (magenta line), n=38 (orange line) and n=34 (blue line) fields of view, each 206 x 206 $\mu m^2$ across on the epithelial layer of n=3 infected bladder-chips. In each case, the protocol results in less than 1 focus of infection per epithelial cell at the end of the 90-minute infection period. The dotted lines and the shaded regions represent the standard deviation.

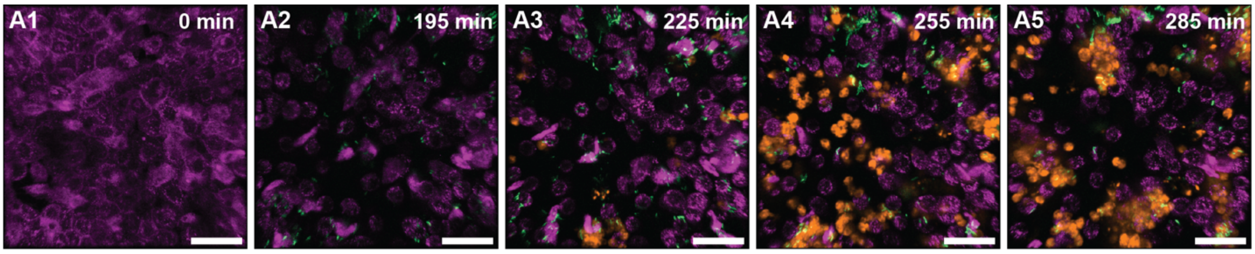


**Figure S5: Timeseries highlighting neutrophil diapedesis and swarm formation.** Additional images that highlight the diapedesis of neutrophils across the epithelial-endothelial barrier and the formation of neutrophil swarms. Bladder epithelial cells (magenta) and neutrophils (amber) were identified with membrane (Cell Mask Orange) and cytoplasmic (Cell Tracker Deep Red) dyes, respectively. UPEC identified via constitutive expression of YFP are shown in green. In all images, scale bar = 50 µm.

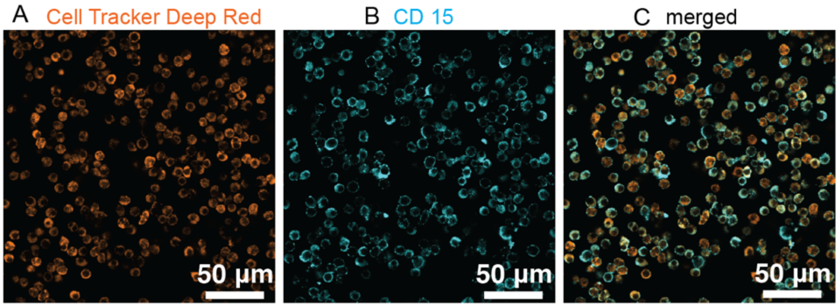


**Figure S6: Neutrophils isolated from human blood are CD15+.**
Images of neutrophils isolated via negative depletion from human blood and labelled with a cytoplasmic dye (Cell Tracker Deep Red) (**A**) and immunostained with an anti-CD15 antibody (**B**). (**C**) Merged image for both channels confirms that all neutrophils are CD15+.

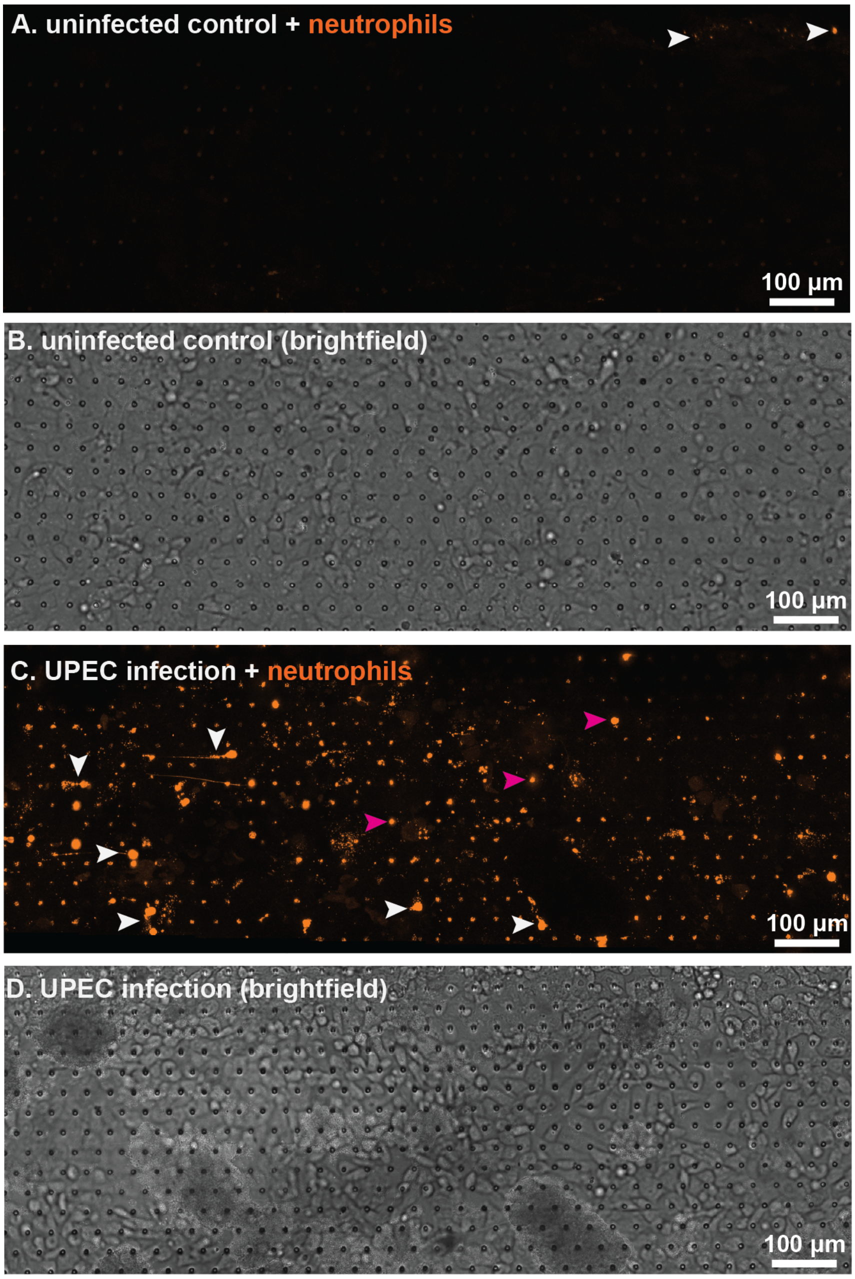


**Figure S7: Neutrophil attachment to endothelial cells is enhanced upon bacterial infection.**

Fluorescent (**A**) and brightfield (**B**) imaging of the endothelial layer of an uninfected bladder-chip. The few neutrophils (identified by CellTracker Deep Red, amber) attached to the endothelial layer are indicated by white arrowheads. Fluorescent (**C**) and brightfield (**D**) imaging of the endothelial layer of an infected bladder-chip, 1.5 hours after infection of the epithelial layer. Neutrophils attached to the endothelial layer are marked by white arrowheads and examples of diapedesis though the PDMS pores to the epithelial layer are marked with magenta arrowheads. In all panels, neutrophils (amber) were introduced into the vascular channel of the bladder-chip under a flow rate of 3 ml/hour corresponding to a shear stress $\eta$=1 dyne/cm$^2$.

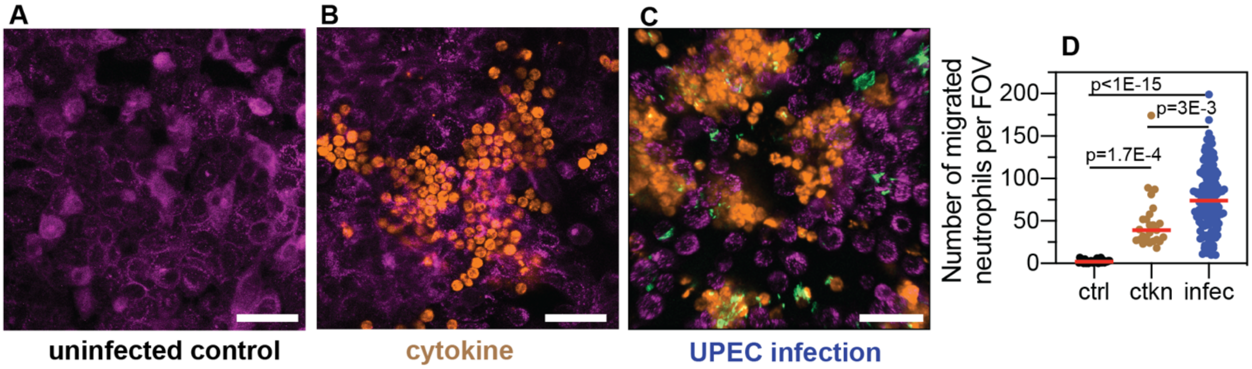


**Figure S8: Neutrophil diapedesis is stimulated by a pro-inflammatory cytokine gradient across the epithelial-endothelial barrier.**

(**A**) Representative images of the epithelial layer of an uninfected control bladder-chip, 2 hours after the introduction of neutrophils in the endothelial channel. No neutrophil diapedesis is observed. (**B**) Representative image of the epithelial layer of an uninfected control bladder-chip exposed to a cocktail of pro-inflammatory cytokines (Interleukin-1α, Interleukin-1β, Interleukin-6 and Interleukin-8, each at 100 ng/ml) added to the diluted urine on the epithelial side and maintained under flow for 2 hours. Epithelial cells (magenta, identified via CellMask Orange) and neutrophils (amber, identified via CellTracker Deep Red) are shown. **C**) Representative image of the epithelial layer of an infected bladder-chip 2 hours after the introduction of neutrophils in the endothelial channel. (**D**) Scatterplot of maximum number of neutrophils detected in 206 x 206 $\mu m^2$ fields of view under control (n= 26), cytokine stimulation (n=26) and infection (n=130). P-values were calculated using Kruskal-Wallis ANOVA Test. Red lines represent median values. Scale bars, 50 μm.

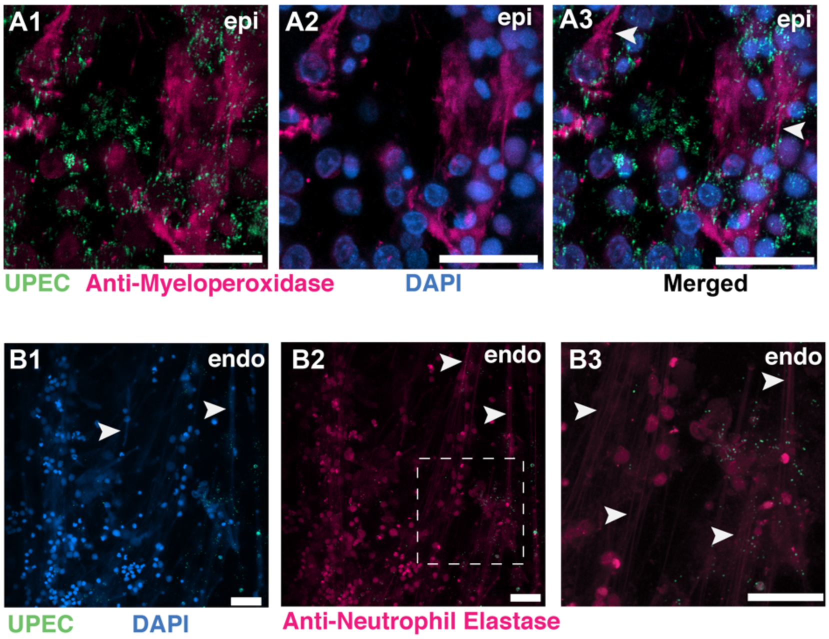


**Figure S9: NETs formation on the epithelial and endothelial layers of an infected bladder-chip.**
(**A1**-**A3**) Additional example of NETs formation by neutrophils on the epithelial layer (epi) of an infected bladder-chip. Neutrophils are identified via immunostaining with an anti-myeloperoxidase antibody (**A1**-**A2**). UPEC identified via YFP expression are shown in spring green (**A1**). Nuclear labelling with DAPI is shown in azure (**A2**). A merged image is shown in **A3**. (**B1**-**B3**) An example of NETs formation by neutrophils on the endothelial layer (endo) of an infected bladder-chip. UPEC identified via YFP expression are shown in spring green and nuclear labelling with DAPI is shown in azure (**B1**). Neutrophils are identified via immunostaining with an anti-neutrophil elastase antibody (**B2**). A zoomed image corresponding to the white dashed box in **B2** is shown in **B3**. In all images, scale bar = 50 µm.

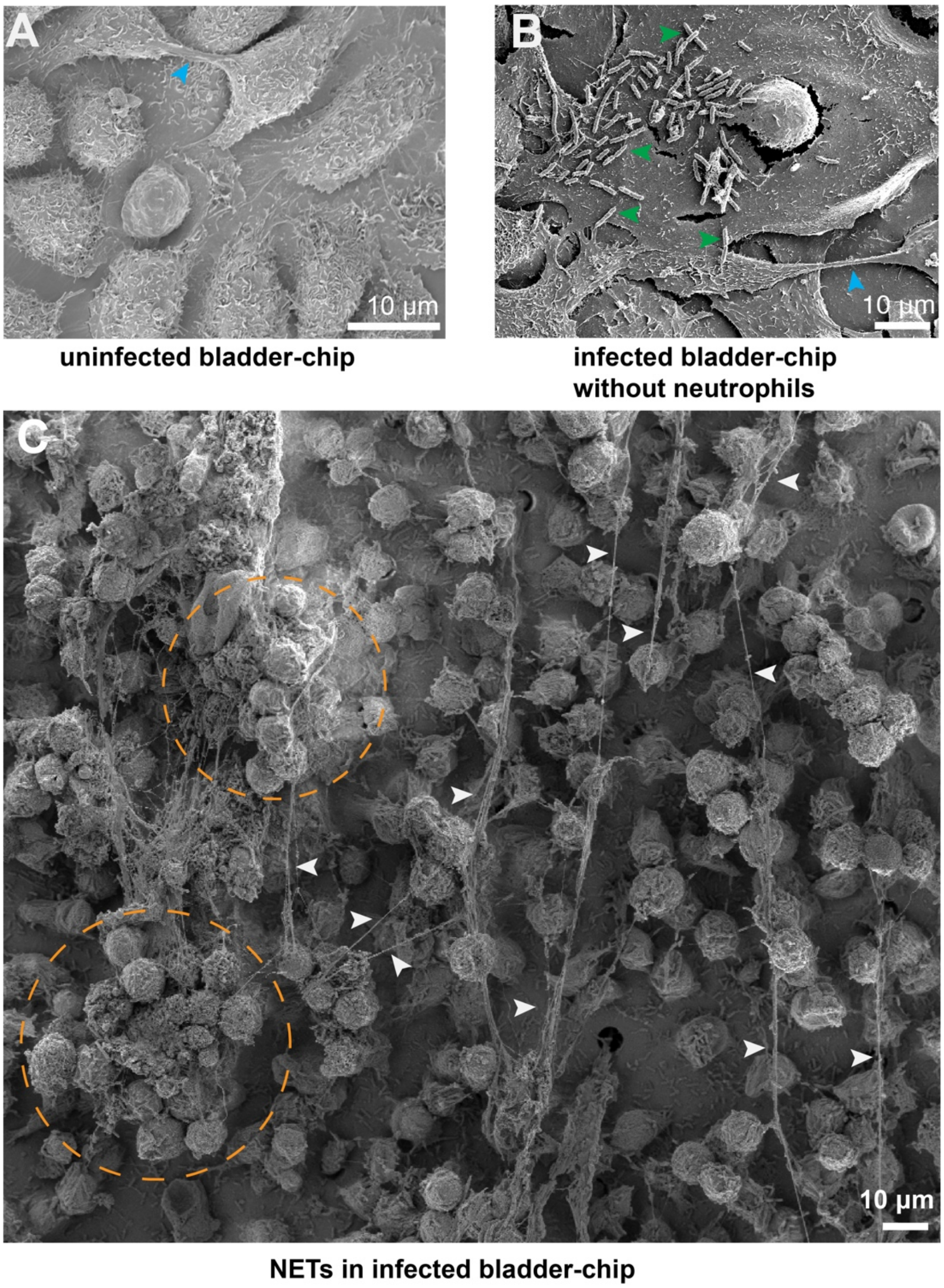


**Figure S10: SEM characterization of uninfected and infected bladder-chips.**

(**A**) SEM image of the confluent epithelial layer of an uninfected bladder-chip. Appendages between epithelial cells are indicated by cyan arrowheads. (**B**) Example from an infected bladder-chip without the addition of neutrophils. Long filaments characteristic of NET formation is not observed. Individual UPEC on the surface of the epithelial cells are indicated by green arrowheads. Appendage between epithelial cells is indicated by a cyan arrowhead (**C**) Additional example of formation of NETS by a large swarm of neutrophils on the epithelial layer of an infected bladder-chip. NETs are indicated by white arrowheads, and large clusters of neutrophils are indicated by dashed amber circles.

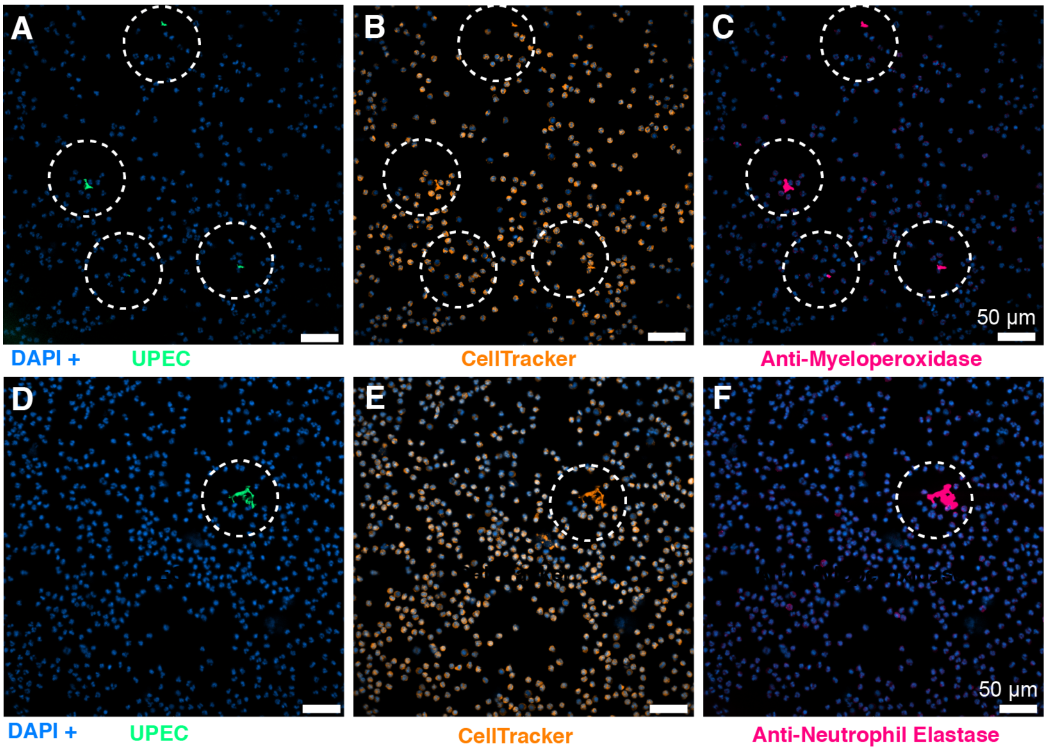


**Figure S11: Neutrophils do not form NETs in response to shear stress in the bladder-chip.**

Neutrophils infused through the vascular channel of an infected bladder-chip were collected and characterized via immunofluorescence for myeloperoxidase (**A-C**) and neutrophil elastase expression (**D-F**) to identify the formation of NETs, indicated by dotted white circles. All neutrophils are labelled by the cytoplasmic CellTracker dye (shown in amber in **B**, **E**). Both myeloperoxidase expression (marked with dotted white circles in **C**) and elastase expression (marked with dotted white circles in **F**) coincide with infected neutrophils (**A**, **D**). UPEC are identified via YFP expression and colored spring green, nuclear labelling is indicated in azure. Scale bars, 50 µm in all panels.

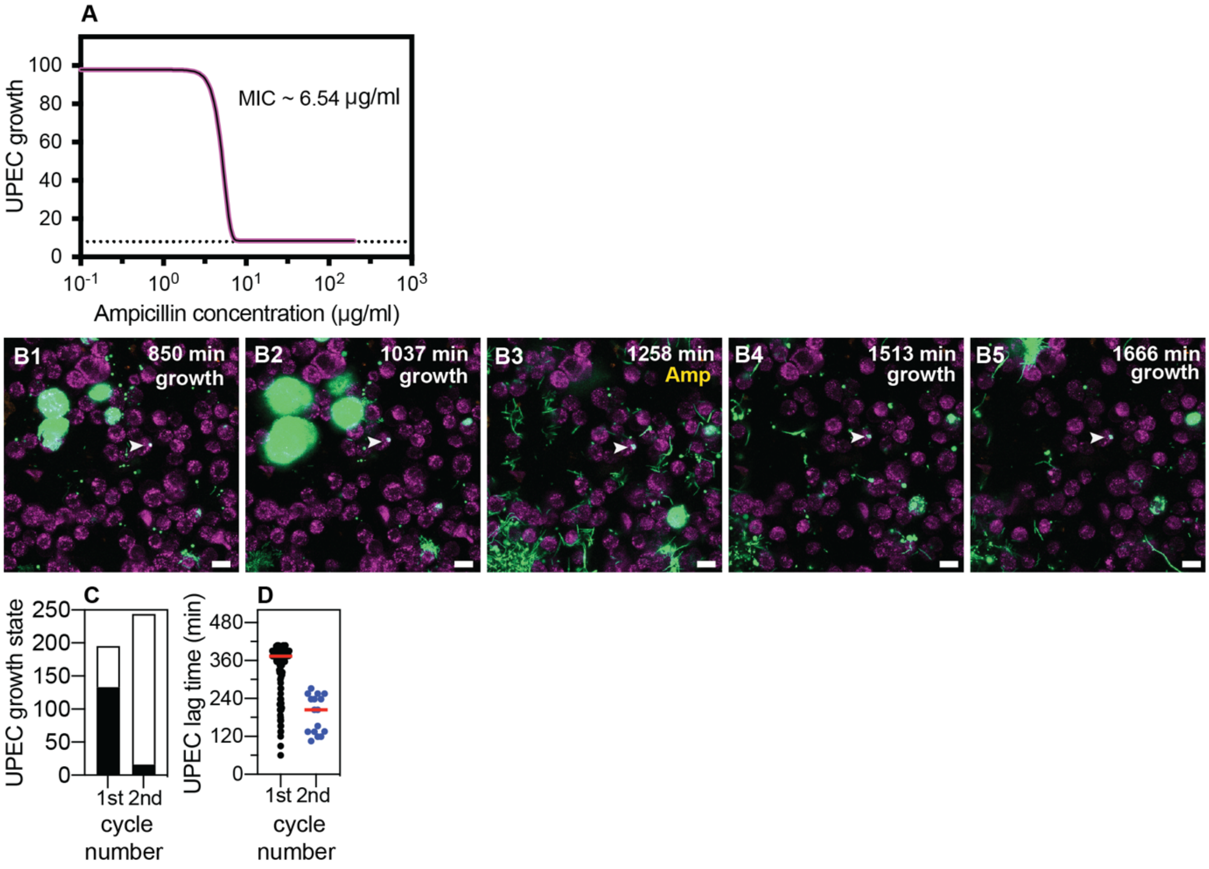


**Figure S12: Non growing UPEC in response to ampicillin administration in the bladder-chip.**

(**A**) Measurement of ampicillin minimum inhibitory concentration (MIC) in endothelial cell medium for the UPEC strain used in these experiments. (**B1**-**B5**) Example of a non-growing clump of UPEC within an epithelial cell following the first growth cycle (indicated by white arrowheads in all images). The bacteria are non-growing throughout the first growth cycle (**B1**, **B2**), the second cycle of ampicillin treatment (**B3**) and subsequently after the removal of antibiotic (**B4**, **B5**). (**C**) Classification of the growth state of intracellular bacterial microcolonies as growing (black) or non-growing (white) across n=108 fields of view in total from n=3 infected bladder chips during the first and second growth periods. (**D**) Scatter plot for the distribution of lag time (measured as the time taken to resume growth after removal of antibiotic) for intracellular bacterial microcolonies during the first (n=133) and second (n=16) growth cycles. Red line represents the median value. Scale bars, 10 μm in **B1**-**B5**.

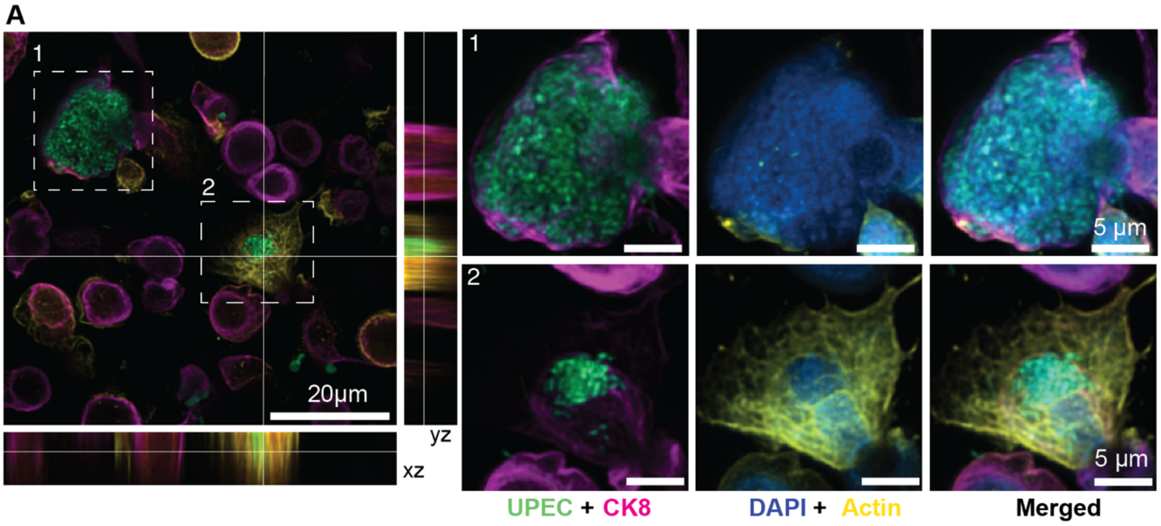


**Figure S13: Immunofluorescence characterization of IBC formation.**
(**A**) Confocal images of two IBCs within epithelial cells with different morphologies. UPEC are labelled in spring green, anti-CK8 staining is shown in magenta, F-actin labelling is shown in yellow, and nuclear labelling with DAPI is shown in azure. IBCs on the infected bladder chip were fixed 13 hours after UPEC infection and 6 hours into the 1st growth cycle.

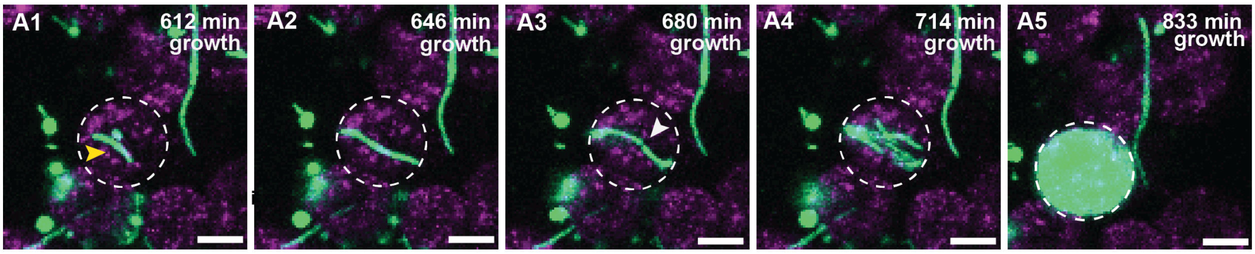


**Figure S14: IBC formation from filamentous UPEC.**
(**A1**-**A5**) Additional example of intracellular growth of filamentous UPEC that develops into an IBC. The growing filament (**A1**-**A2**) is eventually restricted by the cellular volume and bends (**A3**) before reductive division (marked by white arrowhead in **A3**) and IBC formation occurs. Scale bars, 10 μm in **A1**-**A5**.

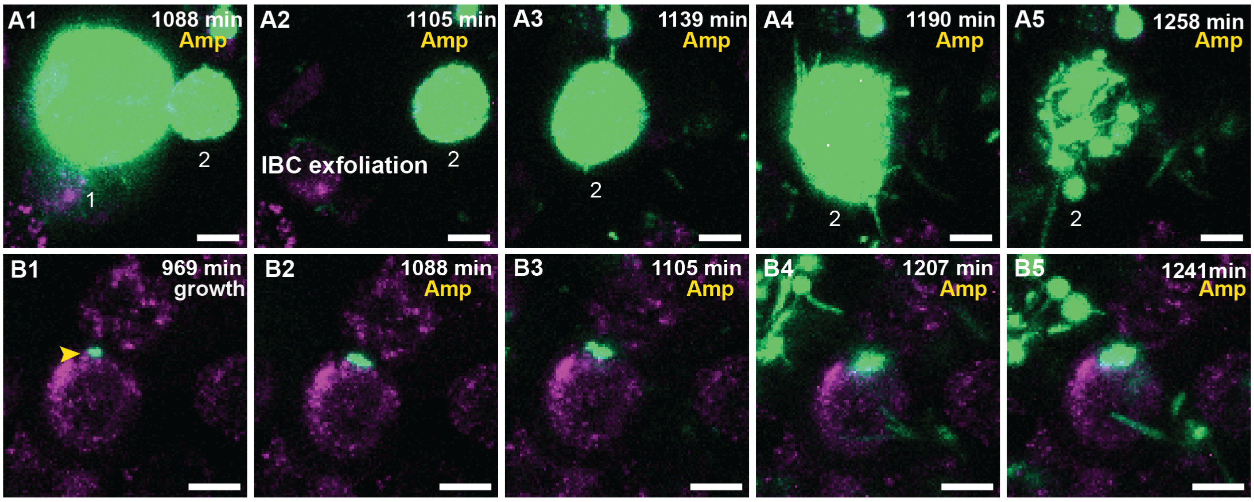


**Figure S15: UPEC growth in IBCs during ampicillin treatment**
(**A1**-**A5**) Additional examples of UPEC growth during ampicillin treatment. (**A1**) Two IBCs (marked 1 and 2) at the start of ampicillin treatment. The IBC marked 1 exfoliates (**A2**) whereas the remaining IBC-2 continues to grow (**A1**-**A3**). Towards the end of this period, the bacteria filament (**A4**) before killing due to the antibiotic is observed (**A5**). (**B1**-**B5**) High resolution time-series that highlights bacterial growth within an IBC prior to (**B1**) and during (**B2-B5**) administration of ampicillin. The bacterial volume within this IBC is not diminished by antibiotic treatment (**B5**). Scale bars, 10 µm in all the panels.

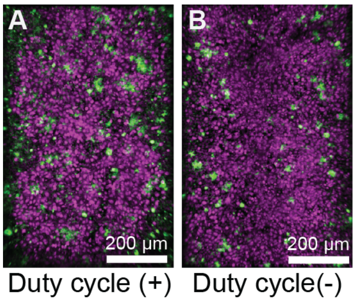


**Figure S16: UPEC infection leads to a higher bacterial burden in bladder-chips perturbed with duty cycle.**
Additional examples of UPEC growth in infected bladder chips with (**A**) and without (**B**) duty cycle.

### Supplementary movie 1

File Name: **SMov1** Description: Infection of bladder-chip with UPEC, diapedesis of neutrophils across the epithelial-endothelial barrier and formation of swarms **(**Fig 2: B1-B5**)**.
Stage 1: Prior to infection, the uninfected cells of the epithelial layer (magenta) are imaged (0-120 mins).
Stage 2: UPEC (green) are introduced into the epithelial channel via flow and infection proceeds under flow for 120-210 min.
Stage 3: Neutrophils (amber) are introduced into the vascular channel of the infected bladder-chip via flow (210 min onwards).
Stage 4: Neutrophils undergo diapedesis within 15-30 minutes and are visible on the epithelial side (210-240 min). UPEC can be seen internalized by neutrophils.
Stage 5: Neutrophils aggregate and form a neutrophil swarm (240 min onwards) on the epithelial side.

### Supplementary movie 2

File Name: **SMov2**, Description: Neutrophil diapedesis and swarm formation on the epithelial side of UPEC infection **(**Fig S5: A1-A5**)**.
Stage 1: Prior to infection, the uninfected cells of the epithelial layer (magenta) are imaged (0-120 mins).
Stage 2: UPEC (green) are introduced into the epithelial channel via flow and infection proceeds under flow for 120-210 min.
Stage 3: Neutrophils (amber) are introduced into the vascular channel of the infected bladder-chip via flow (210 min onwards).
Stage 4: Neutrophils undergo diapedesis within 15 minutes and are visible on the epithelial side (210-240 min). UPEC can be seen internalized by neutrophils.
Stage 5: Neutrophils aggregate and form a neutrophil swarm (240 min onwards) on the epithelial side.

### Supplementary movie 3

File Name: **SMov3**, Description: Formation of intracellular bacterial community inside epithelial cell arising from few bacteria **(**Fig 3: B1-B5).
An intracellular bacterial community is seeded inside a bladder epithelial cell (magenta, stained with Cell Mask Orange) by few UPEC (green) to form an intracellular bacterial community during the first growth cycle (578-1054 min, ca. 8 hr.). During subsequent ampicillin treatment (1071-1258 min, ca. 3 hr.), the IBC shed bacteria and eventually exfoliated from the epithelial layer.

### Supplementary movie 4

File Name: **SMov4**, Description: IBC shedding and exfoliation **(**Fig 3: G1-G5).
UPEC (green) divide and proliferate intracellularly in three epithelial cells (magenta, stained with Cell Mask Orange) to form IBCs. Two late-stage IBCs shed bacteria (ca. 935-969 min). One of these two IBCs subsequently exfoliated (ca. 986 min) whereas the other IBC shrank in volume due to the loss of shed bacteria. The third IBC did not shed bacteria during the first growth cycle (578-1054 min, ca. 8 hr.).

### Supplementary movie 5

File Name: **SMov5**, Description: Filamentous bacterial growth within an IBC (Fig 3:H1-H5).
Filamentous UPEC (green) grow intracellularly in an epithelial cell (magenta, stained with Cell Mask Orange) to form an IBC during the second growth cycle (1275 min onwards). Shedding

of bacteria, some of which appear filamentous was subsequently observed later in the time series (ca. 1649-1768 min).

#### Supplementary movie 6

File Name: **SMov6**, Description: Filamentous bacterial growth within an IBC **(**Fig S14:A1-A5).

Filamentous UPEC (green) grow intracellularly in an epithelial cell (magenta, stained with Cell Mask Orange) to form an IBC during the first growth cycle (578-1054 min, ca. 8 hr.). The IBC eventually exfoliates during the time series.

#### Supplementary movie 7

File Name: **SMov7**, Description: Bacteria within an IBC can persist and grow within an IBC despite the antibiotic treatment **(**Fig 4: A1-A5).

Proliferation of UPEC (green) within four epithelial cells (magenta, stained with Cell Mask Orange) during the 1st growth cycle (578-1054 min, ca. 8 hr.) leads to the formation of IBCs. The bacteria within the four IBCs persist during the (~40x MIC) ampicillin treatment (1071-1258 min, ca. 3 hr.) In each case, bacterial killing was observed during the ampicillin treatment, but all four IBCs persisted over the course of the treatment. Bacterial growth subsequently resumed within all four IBCs post ampicillin washout during the second growth cycle (1275 min onwards). Two new IBCs were formed during the second growth cycle (1275 min onwards). IBC fluxing and filamentation could be observed towards the end of time series.

#### Supplementary movie 8

File Name: **SMov8**, Description**:** Ampicillin mediated bacterial killing is delayed within an IBC **(**Fig 4: F1-F5**)**.

Proliferation of UPEC (green) within an epithelial cell (magenta, stained with Cell Mask Orange) during the 1st growth cycle (578-1054 min, ca. 8 hr.) to form an IBC. The bacteria within the IBC persisted during the (~40x MIC) ampicillin treatment (1071-1258 min, ca. 3 hr.). Bacterial proliferation continued during the first two hours of the ampicillin treatment (1071-1190 min, ca. 2 hr.). Bacterial killing was subsequently observed later during the ampicillin treatment (1190-1258, ca. 1 hr.). Some bacteria within the IBC persisted throughout the ampicillin treatment and resumed proliferation during the second growth cycle (1275 min onwards).

#### Supplementary movie 9

File Name: **SMov9**, Description: Ampicillin mediated bacterial killing is delayed within an IBC (Fig S15:A1-A5).

Proliferation of UPEC (green) within two epithelial cells prior to the ampicillin treatment during the 1st growth cycle (578-1054 min, ca. 8 hr.). IBC#1 exfoliates from the epithelial layer (ca. 1105 min). Bacteria within IBC#2 continued to proliferate during the first two hours of the ampicillin treatment (1071-1190 min, ca. 2 hr.). Bacterial killing was subsequently observed later during the ampicillin treatment (1190-1258, ca. 1 hr.). IBC#2 eventually exfoliated from the epithelial layer towards the end of the time series (1292 min).

#### Supplementary movie 10

File Name: **SMov10**, Description: Bacteria can continue growing within an IBC for the entire duration of ampicillin treatment (Fig 4: G1-G5).

Proliferation of UPEC (green) within an epithelial cell (magenta, stained with Cell Mask Orange) during the 1st growth cycle (578-1054 min, ca. 8 hr.) to form an IBC. Bacterial growth within the intact IBC continued both during the ampicillin treatment (~40x MIC, 1071-1258 min, ca. 3 hr.) as well as after the ampicillin was washed out during the 2nd growth cycle (1275 min onwards). Eventually, the IBCs shed bacteria (1700-1751 min, ca. 1hr) towards the end of time series.

#### Supplementary movie 11

File Name: **SMov11**, Description: Bacteria can continue growing within an IBC during the ampicillin treatment **(**Fig S15:B1-B5).

Proliferation of UPEC (green) within an epithelial cell (magenta, stained with Cell Mask Orange) during the 1st growth cycle (578-1054 min, ca. 8 hr.) to form an IBC. Bacterial growth within the intact IBC continued both during the ampicillin treatment (~40x MIC, 1071-1258 min, ca. 3 hr) as well as after the ampicillin was washed out during the 2nd growth cycle (1275 min onwards). The late-stage IBC eventually shed bacteria and filamentous bacterial growth (1632-1717 min, ca. 1.5 hr) was also observed towards the end of time series.
